# *Listeria monocytogenes* utilizes glutathione and limited inorganic sulfur compounds as a source of essential L-cysteine

**DOI:** 10.1101/2023.10.16.562582

**Authors:** John C. Berude, Paul Kennouche, Michelle L. Reniere, Daniel A. Portnoy

## Abstract

*Listeria monocytogenes* (*Lm*) is a Gram-positive facultative intracellular pathogen that leads a biphasic lifecycle, transitioning its metabolism and selectively inducing virulence genes when it encounters mammalian hosts. Virulence gene expression is controlled by the master virulence regulator PrfA, which is allosterically activated by host- and bacterially-derived glutathione (GSH). The amino acid L-cysteine is the rate-limiting substrate for GSH synthesis in bacteria and is essential for bacterial growth. Unlike many bacteria, *Lm* is auxotrophic for L-cysteine and must import exogenous cysteine for growth and virulence. GSH is enriched in the host cytoplasm, and previous work suggests that *Lm* utilizes exogenous GSH for PrfA activation. Despite these observations, the import mechanism(s) for GSH remains elusive. Analysis of known GSH importers predicted a homologous importer in *Lm* comprised of the Ctp ABC transporter and the OppDF ATPases of the Opp oligopeptide importer. Here, we demonstrated that the Ctp complex is a high-affinity GSH/GSSG importer that is required for *Lm* growth at physiologically relevant concentrations. Further, we demonstrated that OppDF are required for GSH/GSSG import in an Opp-independent manner. These data support a model where Ctp and OppDF form a unique complex for GSH/GSSG import that supports growth and pathogenesis. Additionally, we show that *Lm* utilizes the inorganic sulfur sources thiosulfate and H_2_S for growth in a CysK-dependent manner in the absence of other L-cysteine sources. These findings suggest a pathoadaptive role for partial cysteine auxotrophy in *Lm*, where locally high GSH/GSSG or inorganic sulfur concentrations may signal arrival to distinct host niches.

## Introduction

*Listeria monocytogenes* (*Lm*) is a Gram-positive food-borne pathogen and the causative agent of listeriosis in humans and other mammals (1, 2). This saprophytic bacterium grows ubiquitously in soil and typically encounters mammalian hosts through contaminated food products (3). *Lm* is facultative intracellular pathogen, and bacteria invade host epithelial cells in the gut upon ingestion. Invading bacteria traverse the intestinal epithelium and can cause systemic disease in immunocompromised individuals, such pregnant or elderly individuals (4). During infection, *Lm* must adapt from a saprophytic lifestyle to an intracellular one, shifting its metabolism to better colonize the host intracellular niche. Thus, two factors are central to *Lm* pathogenicity: the ability to utilize host nutrients for intracellular growth and the expression of virulence factors to facilitate pathogenesis.

The master virulence regulator PrfA is essential for *Lm* pathogenesis, as it controls expression of virulence factors necessary for intracellular growth and cell-to-cell spread (5). PrfA is a member of the CRP family of transcriptional regulators that are canonically regulated by allosteric binding of small molecules. Work by Reniere et al and others found that PrfA activity is allosterically controlled by the binding of the low molecular weight thiol glutathione (GSH) (6, 7). GSH is a tripeptide, comprised of glycine, glutamine, and cysteine, and the canonical roles of GSH in cells is to maintain a cytoplasmic reducing environment and detoxify reactive oxygen and electrophilic species (8, 9). Because of its importance in cellular physiology, GSH is produced by all eukaryotic cells and Gram-negative bacteria. Notably, GSH is abundant in the cytoplasm of mammalian host cells where it is present at 1-3mM in most cells and up to 10mM in hepatocytes (9, 10). While most Gram-positive bacteria produce alternative low molecular weight thiols, *Lm* is unusual in that it is one the few Gram-positive species that utilizes GSH as a cytosolic redox buffer and can produce GSH through the bifunctional glutathione synthase, GshF. Bacterial GSH production is essential for full PrfA activation, though previous work has implied that *Lm* can also utilize exogenous GSH for PrfA activation (6, 11).

The amino acid L-cysteine is the rate-limiting component for GSH synthesis and is essential for bacterial growth, where it serves as an amino acid for protein synthesis, an essential component of cytosolic redox buffering systems, and a source of inorganic sulfur for iron-sulfur cluster biogenesis (9). Common sources of nutritional cysteine include the oxidized cysteine species L-cystine, cysteine-containing peptides, GSH and its oxidized counterpart GSSG, L-methionine, and inorganic sulfur sources like sulfate. To obtain L-cysteine from the environment, bacteria have evolved numerous mechanisms for cysteine acquisition and biosynthesis. In *Lm*, L-cystine is imported via the ABC transporter TcyKLMN, which is regulated by the master regulator of the intracellular cysteine pool, CymR, and its regulatory partner, CysK (12). Cysteine-containing peptides are imported by the ABC oligopeptide transporter OppABCDF and the peptide signature of the habitat is speculated to be important for intracellular growth and proper spatiotemporal control of PrfA activation in the cytoplasm of host cells (13).

Reduced L-cysteine is not thought to be a major source of nutritional cysteine in oxic environments because of its high reactivity in aerobic environments and ability to engage in damaging Fenton chemistry (14, 15). Because of this chemistry, intracellular cysteine concentrations are maintained at very low levels and there are few known specific transporters for reduced L-cysteine, with most uptake ascribed to non-specific import via polar amino acid importers (15–17). In *Lm*, reduced L-cysteine has been proposed to be imported by the Ctp ABC transporter, which is comprised of the substrate-binding protein CtaP and the permeases CtpP1 and CtpP2 (18, 19). Considering that the Ctp complex is annotated as an oligopeptide import system, it remains unclear if L-cysteine is the primary substrate for this complex or if import is adventitious.

In addition to importing nutritive cysteine-containing substrates, bacteria can also synthesize L-cysteine from inorganic sulfur sources and L-methionine. Inorganic sulfur sources such as sulfate are commonly imported and converted to H_2_S via the sulfate assimilation pathway, and H_2_S and thiosulfate serve as the primary sulfur sources for L-cysteine biosynthesis via the canonical CysK-dependent two-step biosynthetic pathway (20). While *Lm* has an intact L-cysteine biosynthetic pathway, it lacks the genes for sulfate assimilation and is unable to grow on sulfate and has not been observed to grow on thiosulfate (21–23). L-methionine is the second of the two sulfur-containing amino acids, and it is commonly converted to L-cysteine via the transsulfuration pathway. However, some strains of *Lm* lack the genes for transsulfuration and are not able to grow on L-methionine as a sole sulfur source, including 10403S (23).

Despite the above observations, nothing is known about the mechanisms for exogenous GSH import and utilization or the function of the intact L-cysteine biosynthetic pathway in *Lm*. GSH and GSSG are the most abundant sources of L-cysteine in the host cytosol, which is the preferred host niche for *Lm*, and other pathogenic bacteria such as *E.* coli, *F. tularensis*, and *S. mutans* are known to utilize host GSH for growth and virulence by either importing these molecules directly or by degrading them via the action of secreted γ-glutamyl transpeptidases (24–27). Previous work has suggested that *Lm* is able to import GSH for PrfA activation, as treatment with exogenous GSH is sufficient to activate PrfA in a *gshF* mutant (11). These observations suggest that the locally high GSH concentrations in the host cytosol could be a cue for bacterial arrival to the intracellular niche. Similarly, H_2_S and thiosulfate are present at relatively high concentrations in the lumen of the intestine (1-3mM) and could similarly serve as a cue for bacterial arrival to the host environment (28–30).

Here, we report that *Lm* utilizes diverse L-cysteine sources for growth, including GSH, GSSG, and the inorganic sulfur sources H_2_S and thiosulfate. We demonstrate that the Ctp complex and the OppDF ATPases are required for growth on GSH and GSSG *in vitro*, and that the Ctp/OppDF complex is a high-affinity importer of GSH and GSSG that is required for full virulence in a murine IV infection model. Additionally, we show that *Lm* utilizes the inorganic sulfur sources H_2_S and thiosulfate for growth, and that growth on these substrates is dependent on CysK and the canonical two-step biosynthetic pathway. Together, these findings suggest a pathoadaptive role for the partial cysteine auxotrophy in *Lm*, where locally high GSH/GSSG or thiosulfate concentrations may signal arrival to distinct host niches.

## Results

### *Lm* utilizes diverse sulfur sources for growth in vitro, including GSH, GSSG, and thiosulfate

To test the ability of WT *Lm* to grow on different exogenous L-cysteine sources, we performed broth growth assays in chemically defined synthetic media (LSM) containing all amino acids except L-cysteine and supplemented with either L-cystine or another exogenous L-cysteine source (Fig. 1). We observed normal growth of WT bacteria in LSM supplemented with either 0.5mM L-cystine or 1mM L-cysteine. Bacteria grown in LSM lacking all L-cysteine sources (LSM-cys) doubled once, presumably due to the presence of sufficient L-cysteine from overnight growth, but then failed to grow further. Notably, the LSM-cys growth condition included 0.67mM L-methionine and ∼2mM sulfate, confirming previous genetic and experimental evidence that WT 10403S lacks the ability to synthesize L-cysteine from these sources using the reverse transsulfuration and sulfate assimilation pathways, respectively (21–23, 31). We also observed growth on the Arg-Gly-Asp-Cys (RGDC) oligopeptide as a sole L-cysteine source, which is consistent with previous reports (13). Growth kinetics of WT *Lm* on RGDC were slightly altered relative to growth on L-cystine, demonstrating a longer lag phase, comparable log growth rate, and the same saturation density after 24 hours.

**Figure 1.**
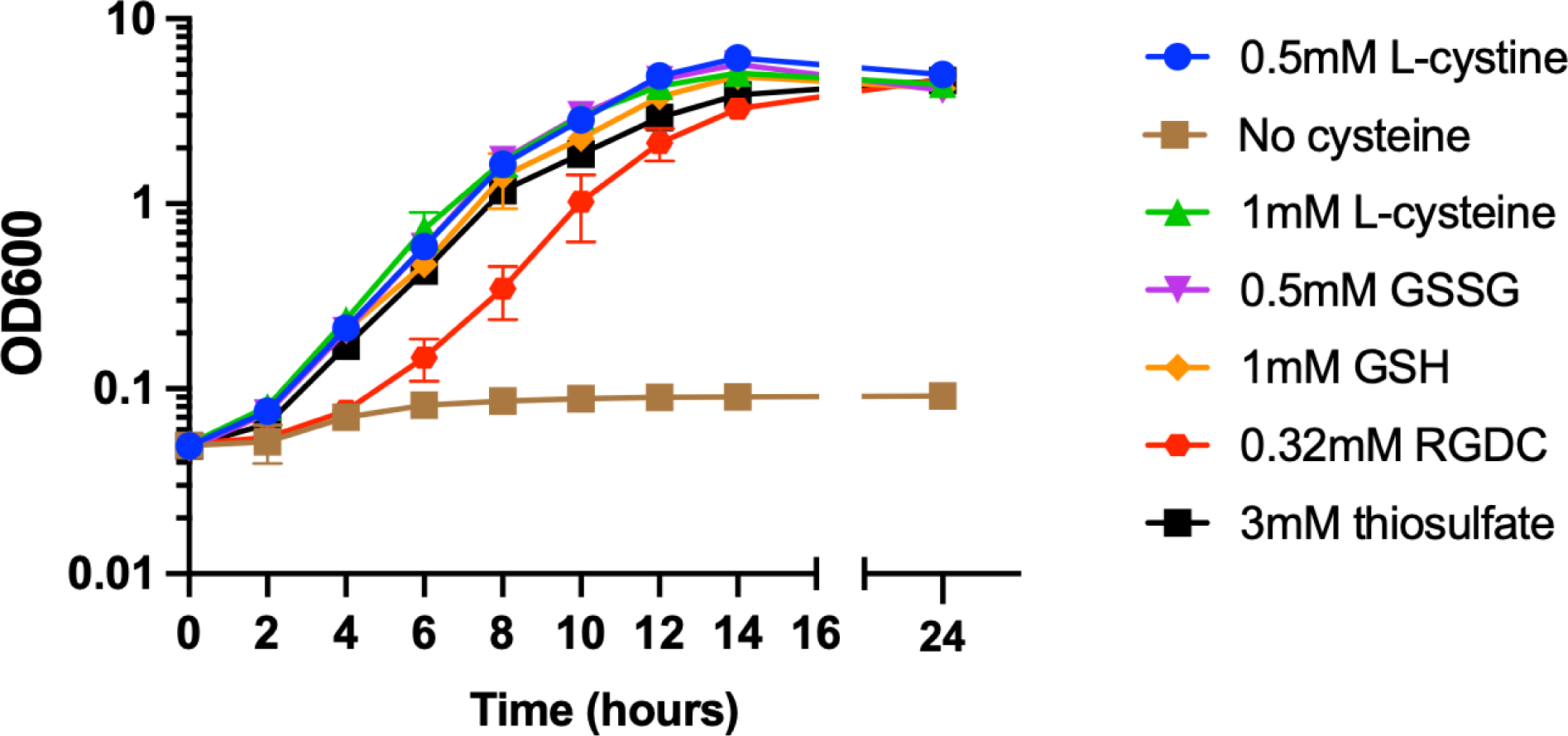
*Lm* utilizes diverse L-cysteine sources for growth *in vitro*. Broth growth of WT *Lm* in cysteine-free media supplemented with the indicated L-cysteine source. Bacteria were grown overnight in LSM supplemented with 0.5mM L-cystine. Cultures were washed once in LSM lacking L-cysteine and inoculated into media supplemented with various physiologically relevant L-cysteine sources. Growth was determined by OD_600_ at the indicated timepoints. Data representative of three independent replicates, error bars represent SD.

While other exogenous L-cysteine sources were used at concentrations that provided 1mM equivalent of L-cysteine, the RGDC peptide was provided at 0.32mM L-cysteine equivalent as previously described due to saturation of the Opp oligopeptide importer (13). While previous studies on PrfA activation have implied that *Lm* can import and utilize exogenous GSH to activate PrfA (6, 11), growth on GSH or its oxidized counterpart, GSSG, as a sole L-cysteine source has not been studied. Here, we found that WT *Lm* utilized both GSH and GSSG for growth in the absence of other L-cysteine sources and that growth kinetics match those of growth on both L-cystine and L-cysteine. Further, we also observed that WT *Lm* grows readily on the inorganic sulfur compound thiosulfate in the absence of other L-cysteine sources, displaying similar growth kinetics to L-cystine, L-cystine, GSH, and GSSG. This observation is contrary to previous observations and suggests that *Lm* is not a complete L-cysteine auxotroph as previously suggested (22, 23, 32).

### H_2_S and thiosulfate utilization requires the enzyme CysK

To better understand the mechanism(s) of thiosulfate utilization in *Lm*, we considered the role of the two-step L-cysteine biosynthetic pathway in facilitating growth on thiosulfate. The L-cysteine two-step biosynthetic pathway is a highly conserved pathway in bacteria that involves the conversion of L-serine to L-cysteine by two enzymes: CysE and CysK (Fig. 2A). In the first step, CysE combines L-serine and acetyl-CoA to form the intermediate O-acetylserine (OAS). In the second step, CysK condenses OAS with sulfide or thiosulfate to form L-cysteine (20). These two genes are in distinct genomic loci in *Lm*, with *cysE* present as part of a cysteine-responsive operon and *cysK* residing at a separate genomic locus (Fig. 2B).

**Figure 2.**
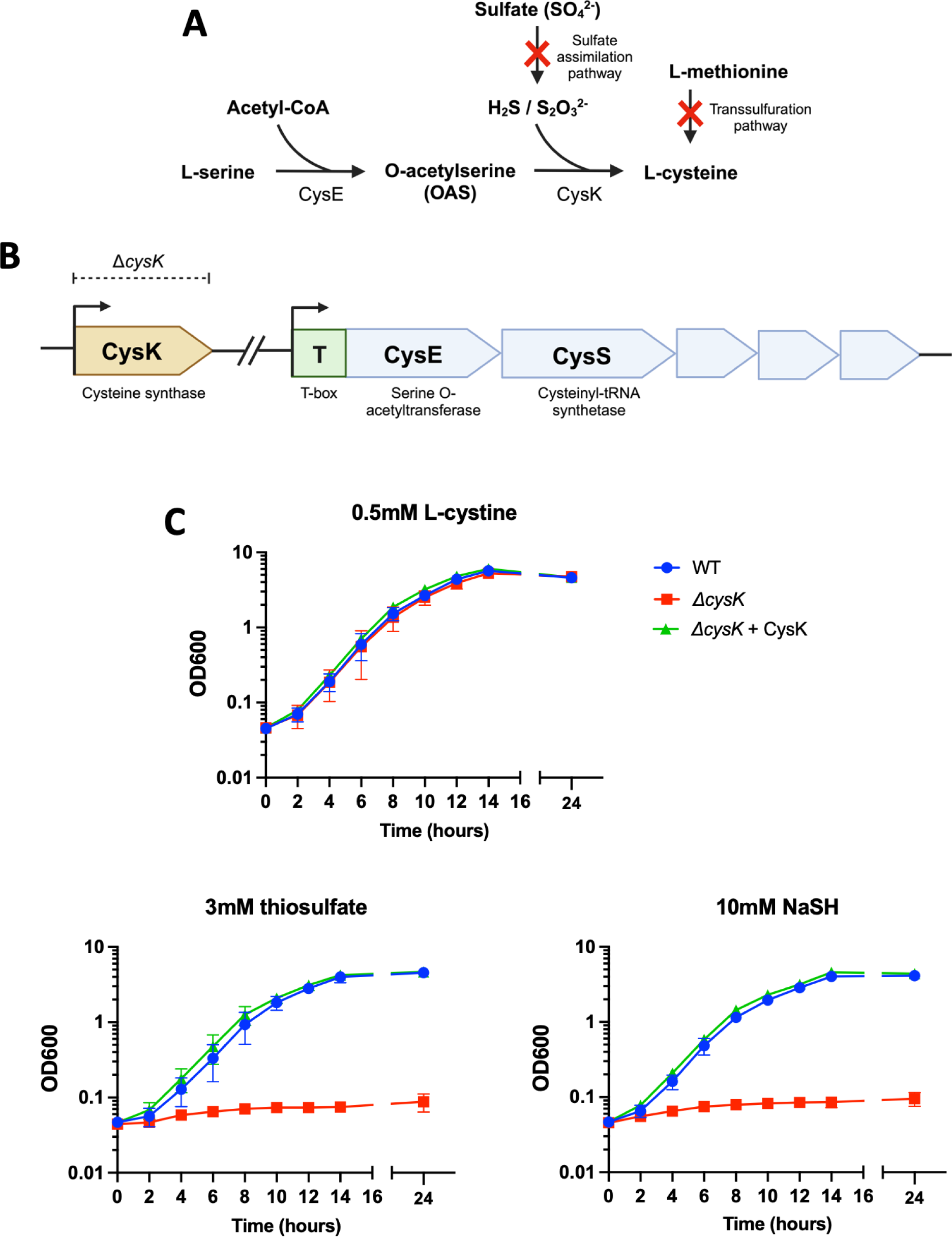
CysK is required for *Lm* growth on thiosulfate and H_2_S. A) Schematic of the two-step L-cysteine biosynthetic pathway in *Lm* from L-serine and inorganic sulfur, including notable non-functional pathways in *Lm*. B) Genomic organization of genes of the two-step biosynthetic pathway. C) Broth growth of the *cysK* mutant and complement in LSM supplemented with either thiosulfate or the H_2_S generator NaSH. Strains were grown overnight in LSM media containing 0.5mM L-cystine, washed once in LSM lacking L-cysteine, and inoculated into media containing either L-cystine or the indicated inorganic sulfur source as the sole sulfur source. Growth was determined by OD_600_ at the indicated timepoints. Data representative of three independent replicates, error bars represent SD.

Typically, sulfide for this pathway is generated by import and reduction of sulfate by the sulfate assimilation pathway in sulfur-reducing bacteria (33), though reduced H_2_S can be acquired from the environment through passive transport across the cell membrane (34). Thiosulfate is generated by the oxidation of H_2_S in aerobic environments and is typically imported by members of the sulfate assimilation pathway or alternative thiosulfate-specific and inorganic ion importers (35). Previous genetic and experimental evidence has shown that WT 10403S *Lm* is unable to utilize exogenous sulfate or thiosulfate to produce L-cysteine, however it has been noted that WT *Lm* harbors an intact CysE/CysK two-step biosynthetic pathway (12, 21–23).

To test the role of the CysE/CysK two-step biosynthetic pathway, we generated a *ΔcysK* mutant and tested its growth on thiosulfate and H_2_S in the absence of other L-cysteine sources (Fig. 2C). Neither the Δ*cysK* mutant nor the complemented strain displayed a defect when grown on L-cystine as a sole L-cysteine source. However, we observed that the Δ*cysK* mutant did not grow on either 3mM thiosulfate or 10mM sodium hydrosulfide (NaSH), which is a potent H_2_S donor. These defects were rescued in both conditions by the complementation with the native *cysK* gene, demonstrating that the CysE/CysK two-step biosynthetic pathway is functional in *Lm* and enables growth on a limited number of inorganic sulfur sources. The concentrations of thiosulfate and NaSH used represent the lowest concentrations that yielded similar growth kinetics in WT *Lm* to the 0.5mM L-cystine control condition (J.C. Berude and D.A. Portnoy, unpublished data). Notably, 10mM NaSH was necessary to achieve similar growth kinetics of WT *Lm* as with 3mM thiosulfate or 0.5mM L-cystine, likely due to the off-gassing of H_2_S generated by the hydrosulfide ion in solution.

Next, we sought to determine if the CysE/CysK biosynthetic pathway plays a role in virulence gene activation in *Lm* during infection. Human tissues are known to produce micromolar quantities of H_2_S (36, 37), however concentrations of up to 1-3mM are present in the anoxic lumen of the intestine (30). The H_2_S concentration in the gut is uniquely high and is toxic to colonic epithelial cells, and thus H_2_S is rapidly converted to thiosulfate in the oxic environment of the outer intestinal lumen (28, 30). This reaction generates similarly high concentrations of thiosulfate (1-3mM) and suggests that these substrates might support *Lm* growth and virulence in the gut. We observed no detectable defect in virulence of the Δ*cysK* mutant in a murine intravenous infection model (Fig. S1), which is a measure of intracellular growth and virulence *in vivo*. We also did not observe a defect in growth of the *cysK* mutant in a murine oral competition assay, in which mice were orally infected with a 1:1 ratio of erythromycin (Erm)-sensitive WT and Erm-resistant *cysK*::Tn bacteria in a Δ*hly* background, which prevents intracellular infection and restricts bacteria to extracellular growth in the lumen of the intestine (Fig. S2). Both WT and *cysK*::Tn Erm-resistant bacteria displayed a slight and equivalent defect in growth relative to Erm-sensitive WT bacteria, which suggests that the slightly low ratio of *cysK*::Tn/WT is due to lower plating efficiency of the Erm resistant strain and is not related to mutation of the *cysK* gene (38). Together, these observations suggest that the CysE/CysK two-step pathway does not play a significant role in growth and virulence of *Lm* in the diverse nutritional landscape of the intestinal lumen.

### The Ctp/OppDF complex imports GSH and GSSG

Next, we sought to elucidate the mechanism underlying GSH and GSSG import in *Lm*. The GsiABCD ABC transporter is a known GSH-specific importer in *E. coli*, and a BLAST search for these genes identified putative homologs in *Lm* that included the genes of the Ctp ABC transporter and the OppDF ATPases from the Opp complex (Table 1). The Ctp operon consists of three genes: a substrate-binding domain (*ctaP*) and two permeases (*ctpP1* and *ctpP2*) but lacks dedicated ATPase domains (Fig. 3A). The Opp operon encodes a conserved oligopeptide importer that includes a substrate-binding domain (*oppA*), two permeases (*oppB* and *oppC*), and two ATPases (*oppD* and *oppF*) (Fig. 3A). In *Lm*, the Opp complex is important for growth on oligopeptides.

**Figure 3.**
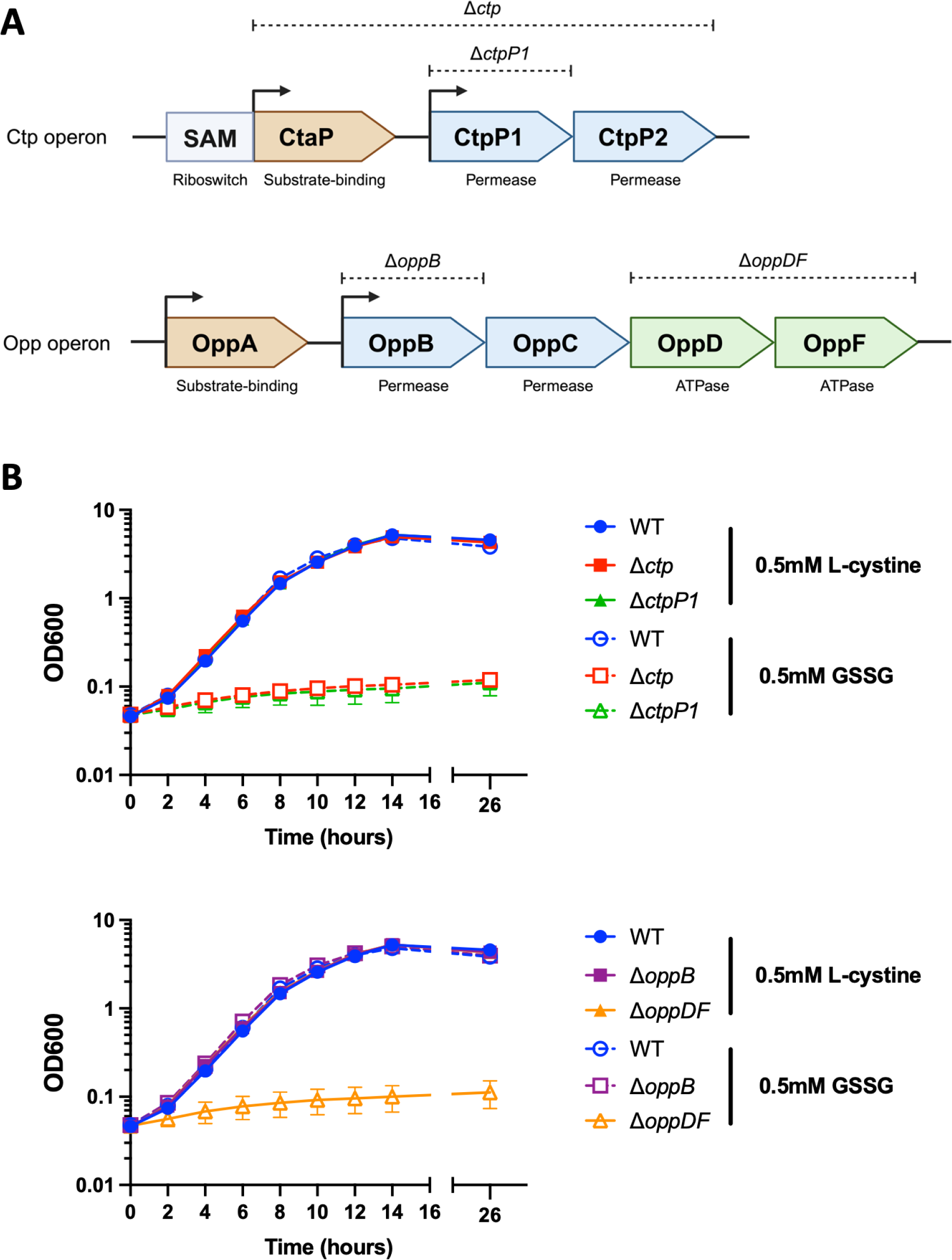
The Ctp operon and OppDF are required for growth on GSSG. A) Genomic organization of genes of the *ctp* and *opp* operons. Dashed lines denote specific mutants constructed. B) Broth growth of the *ctp* and *opp* mutants in LSM supplemented with GSSG. Strains were grown overnight in LSM media containing 0.5mM L-cystine, washed once in LSM lacking L-cysteine, and inoculated into media containing either L-cystine or GSSG as the sole L-cysteine source. Growth was determined by OD_600_ at the indicated timepoints. Data representative of three independent replicates, error bars represent SD.

**Table 1.** Homology of *E. coli* GsiABCD genes to *Lm* Ctp and OppDF genes. Homology determined by BLASTp search. Listed homology scores for the top search hits only.

|  | % homology to *E. coli* GSH ABC importer GsiABCD subunits |  |  |  | Annotated function |
| --- | --- | --- | --- | --- | --- |
| *Lm* genes | GsiA | GsiB | GsiC | GsiD | *Lm* / *E. coli* |
| CtaP (Lmo0135) | - | 26% | - | - | Substrate-binding |
| CtpP1 (Lmo0136) | - | - | 38% | - | Permease |
| CtpP2 (Lmo0137) | - | - | - | 40% | Permease |
| OppD (Lmo2193) | 42% | - | - | - | ATPase |
| OppF (Lmo2192) | 48% | - | - | - | ATPase |

Previous work with the OppDF ATPases has suggested that they may play an important role in PrfA activation and intracellular growth, though it remains unclear if the core Opp complex (OppABC) is also required (13, 39). Additionally, even though the Ctp ABC transporter was initially annotated as a peptide importer in *Lm*, it has been suggested that the complex is involved in high-affinity import of reduced L-cysteine (18, 19).

To determine if the Ctp and Opp complexes are involved in GSH and GSSG import, we generated mutants in both pathways and assessed broth growth in LSM supplemented with either L-cystine or GSSG as a sole L-cysteine source. As expected, we observed normal growth kinetics of all strains (WT, Δ*ctp*, Δ*ctpP1*, Δ*oppB*, Δ*oppDF*) in 0.5mM L-cystine (Fig. 3B). WT *Lm* displayed normal growth kinetics when grown in 0.5mM GSSG, though no growth was observed in the Δ*ctp*, Δ*ctpP1*, and Δ*oppDF* mutants, suggesting that the Ctp complex and OppDF ATPases are both required for growth on GSSG (Fig. 3B). The Δ*oppB* mutant displayed normal growth kinetics on both L-cystine and GSSG, suggesting that the core Opp complex is not necessary for GSSG import (Fig. 3B).

To further elucidate the contributions of the Ctp and Opp complexes for growth on different L-cysteine sources, we tested the growth of mutants and their complements in saturation endpoint growth assays. As previously observed, WT *Lm* displayed normal growth with 0.5mM L-cystine, 1mM L-cysteine, 0.5mM GSSG, 1mM GSH, and 0.32mM RGDC, and failed to grow in LSM lacking any L-cysteine source (Fig. 4A). Mutants in the Ctp complex (Δ*ctp* and Δ*ctpP1*) displayed normal growth on L-cystine, L-cysteine, and RGDC, but were unable to grow on either GSH or GSSG, and growth on these substrates was recovered by complementation (Fig. 4A). This result suggests that the Ctp locus is not required for growth on L-cysteine as previously reported, but instead for growth on GSH and GSSG. Deletion of Δ*oppDF* similarly impaired *Lm* growth on GSH and GSSG but also eliminated growth on RGDC, which is consistent with the established role of the Opp complex for growth on oligopeptides. Growth of the Δ*oppDF* mutant on all three substrates could be recovered by complementation with *oppDF*, though complementation of the Δ*oppDF* mutant with either *oppD* or *oppF* alone did not rescue growth on GSH, GSSG, and RGDC, suggesting that both ATPase subunits are required for the proper function of both the Ctp and Opp complexes (Fig. 4B). As previously observed, the Δ*oppB* mutant displayed no growth defect on either GSH or GSSG, but also exhibited a partial defect when grown on RGDC that was rescued by complementation (Fig. 4C). To further explore the contribution of the Ctp complex to growth on short cysteine-containing peptides, we tested the growth of a Ctp/Opp double mutant. Consistent with observations with the Δ*oppDF* mutant, a Δ*ctpP1*/*oppB*::Tn grew like WT *Lm* on L-cystine and L-cysteine, but was unable to grow on GSH, GSSG, or RGDC (Fig. 4D). Complementation with *oppB* rescued growth on RGDC, but not GSH or GSSG, while complementation with *ctpP1* fully rescued growth on GSH and GSSG and partially rescued growth on RGDC. Consistent with the initial annotation of the Ctp locus as a peptide transporter, these observations suggest that the Ctp complex may also facilitate import of short cysteine-containing peptides.

**Figure 4.**
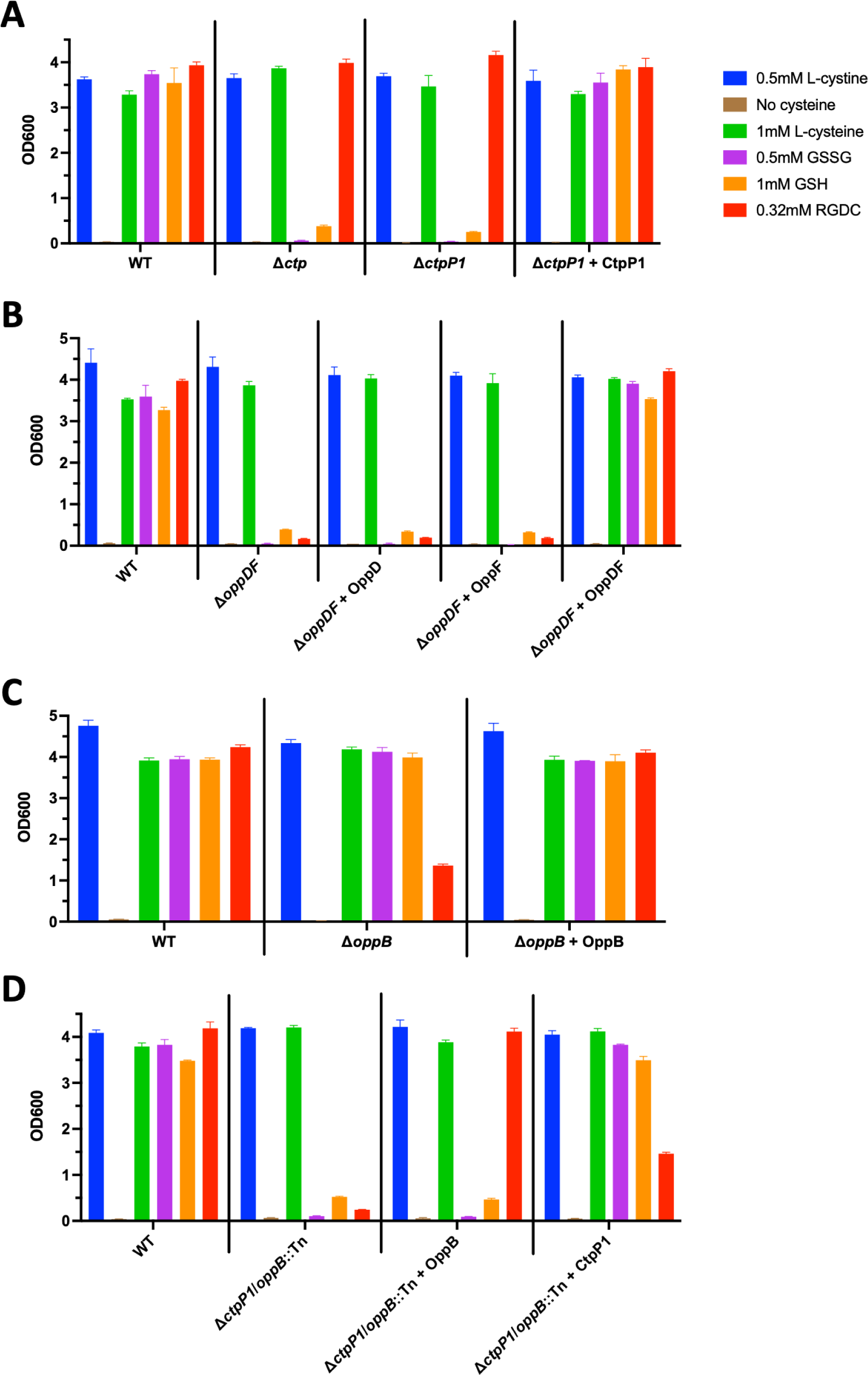
The Ctp operon and OppDF are required for growth on GSH and GSSG. Endpoint saturation broth growth of mutants and complements in cysteine-free LSM media supplemented with the indicated L-cysteine source. A) Broth growth of *ctp* mutants and complement. B) Broth growth of the *oppDF* mutant and complements. C) Broth growth of the *oppB* mutant and complement. D) Broth growth of the *ctpP1*/*oppB*::Tn double mutant and complements. Strains were grown overnight in LSM containing 0.5mM L-cystine, washed once in LSM lacking L-cysteine, and inoculated into media containing the indicated L-cysteine source as the sole L-cysteine source. Growth was determined by OD_600_ at the indicated timepoints. Data representative of three independent replicates, error bars represent SD.

Because LSM is a defined, synthetic medium that represents an artificial bacterial growth environment, we sought to determine if the Ctp complex plays a role in more complex growth conditions *in vitro*. Brain-Heart Infusion (BHI) medium is a rich, complex medium that is commonly used to cultivate fastidious pathogenic microorganisms like *Lm*. The medium derives most of its nutrients from an infusion of host tissues, peptone, and glucose, with L-cysteine presumably present as GSH and/or GSSG, L-cystine, and as part of oligopeptides of various length. We tested the growth of our mutants in BHI and observed growth defects only in the Δ*oppDF* and Δ*ctpP1*/*oppB*::Tn double mutants, and not with the Δ*ctp*, Δ*ctpP1*, and Δ*oppB* single mutants (Fig. S3). This result suggests that *Lm* can acquire sufficient L-cysteine from either GSH/GSSG or oligopeptides, but that at least one of these import pathways is required for full *Lm* growth in BHI. Taken together, these observations provide evidence of a novel GSH/GSSG-specific importer in *Lm* that is comprised of the Ctp ABC transporter complex and the OppDF ATPases.

### The Ctp/OppDF complex is a high-affinity GSH/GSSG importer

To further elucidate the role of the Ctp/OppDF complex in GSH and GSSG import, we tested the affinity of the Ctp complex for both substrates in saturation endpoint growth assays. We observed that WT *Lm* grew on as little at 5uM GSSG, with normal growth observed at concentrations at or above 50uM (Fig. 5A). The Δ*ctp* mutant did not grow at any of the tested GSSG concentrations (1uM-5mM), suggesting that the Ctp complex is a high-affinity GSSG importer. Similarly, we tested *Lm* growth on concentrations of GSH from 1uM-10mM, and we observed that WT *Lm* grows on GSH concentrations as low as 50uM, with normal growth observed at or above 100uM (Fig. 5B). The Δ*ctp* mutant did not grow in GSH concentrations at or below 0.5mM, with slight growth at 1mM, moderate growth at 2mM, and normal growth observed at 3mM GSH and above. Taken together, these data suggest that the Ctp complex is a high-affinity GSH importer, but that there are also alternative mechanisms of GSH acquisition in *Lm*.

**Figure 5.**
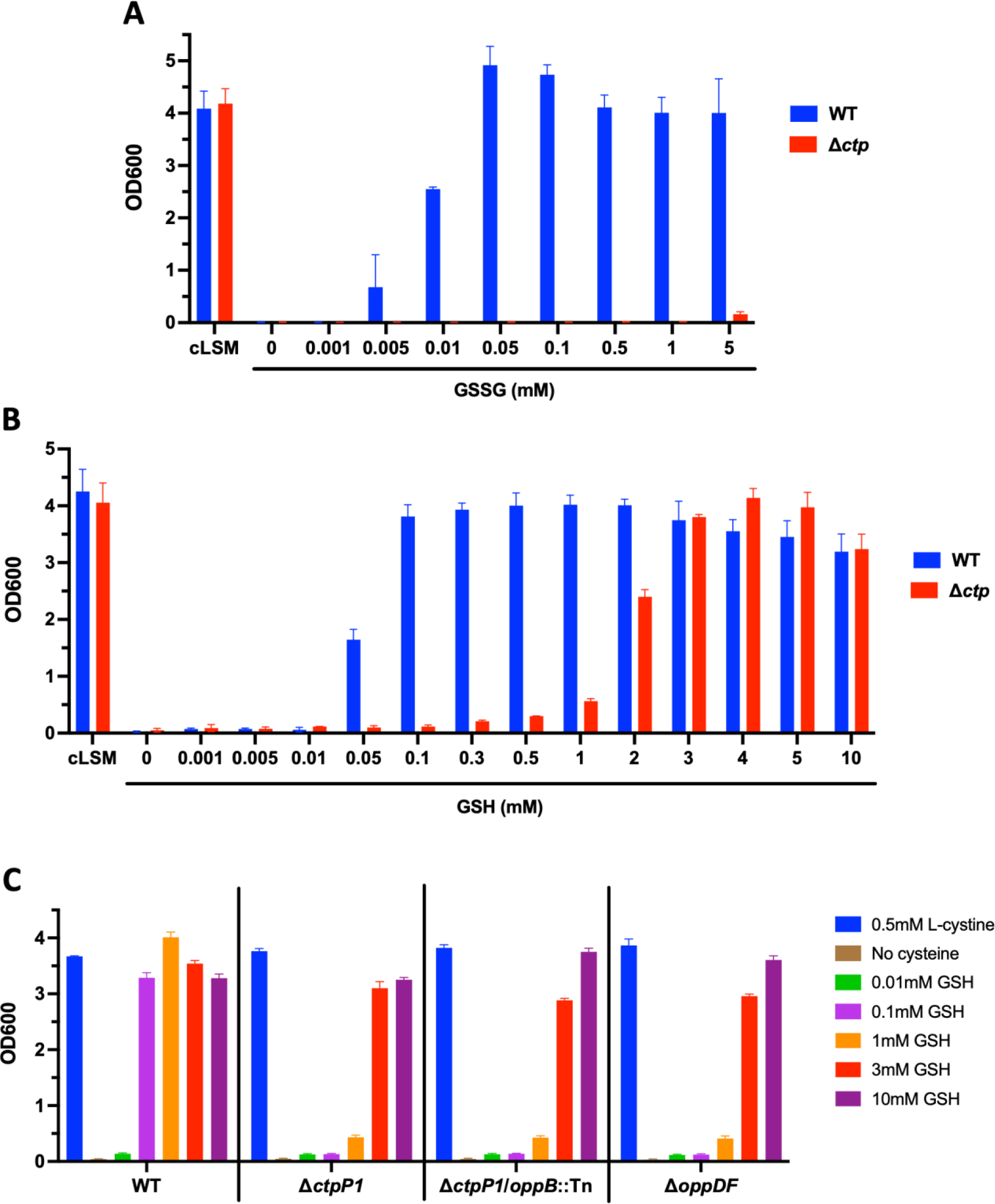
The Ctp complex is a high-affinity GSH/GSSG importer. Endpoint saturation broth growth of mutants and complements in cysteine-free LSM media supplemented with the indicated L-cysteine source. A) Broth growth of the *ctp* mutant in indicated concentrations of GSSG. B) Broth growth of the *ctp* mutant in indicated concentrations of GSH. All GSH media supplemented with 5mM TCEP. C) Broth growth of the *ctpP1*, *oppDF*, and *ctpP1*/*oppB*::Tn mutants on indicated concentrations of GSH. Strains were grown overnight in LSM containing 0.5mM L-cystine, washed once in LSM lacking L-cysteine, and inoculated into media containing the indicated L-cysteine source as the sole L-cysteine source. Growth was determined by OD_600_ at the indicated timepoints. Data representative of three independent replicates, error bars represent SD.

Because GSH is a tripeptide, we considered if the Opp complex may also import GSH and account for Ctp-independent growth on higher concentrations of GSH. Previous experiments with the Δ*ctpP1*/*oppB*::Tn double mutant strain demonstrated that complementation with *oppB* was not sufficient to recover growth on 1mM GSH (Fig. 4D), suggesting that the Opp complex is not able to import GSH. To confirm this hypothesis, we tested growth of Ctp and Opp mutants in varying concentrations of GSH. Neither the ΔctpP1, ΔctpP1/*oppB*::Tn, or Δ*oppDF* mutants were able to grow on GSH concentrations of 1mM or less, and all demonstrated normal growth at 3mM GSH and above, suggesting that the Opp complex is not an alternate GSH importer (Fig. 5C).

Overall, these observations suggest that the Ctp/OppDF complex functions as a high-affinity GSH/GSSG importer in *Lm*, and that there is likely an alternative low-affinity GSH import mechanism that is independent of the Ctp and Opp complexes.

### The Ctp and Opp complexes are required for virulence in a murine IV infection model

GSH is the most abundant source of L-cysteine in the mammalian cytoplasm, and imported GSH likely contributes directly to PrfA activation, virulence gene expression, and intracellular growth. Thus, the Ctp complex is of particular interest as a GSH/GSSG-specific importer and potential modulator of virulence in *Lm*. Previous work has demonstrated that mutants in the Ctp complex exhibit variable defects in a murine IV infection model (18, 19), but it remains unclear if these defects are due to decreased PrfA activation needed for virulence, reduced access to nutritional L-cysteine needed for intracellular growth, or both. We sought to recapitulate these findings using an IV infection model and assess the relative contributions of the Ctp and Opp complexes to PrfA activation and nutrient acquisition *in vivo*.

Contrary to previous findings, pooled data from three independent intravenous infection experiments did not reveal a significant impact on virulence with the Δ*ctp* mutant, suggesting that the Ctp complex alone is dispensable for growth and virulence activation *in vivo*. However, we observed a significant impact on virulence with the Δ*oppDF* mutant, which displayed a ∼150-fold decrease in colony-forming units (CFU) in the livers of infected mice at 48 hours post-infection (Fig. 6). To determine if these defects were due to a reduction in PrfA activation or a defect in nutrient acquisition, we compared the virulence defects of mutants in the WT 10403S background to those in the PrfA* background, which encodes a constitutively active form of PrfA that does not require allosteric binding of GSH for activation (6, 40). The ∼150-fold virulence defect in the Δ*oppDF* mutant was reduced to ∼15-fold in the PrfA* background, which suggests that the Ctp and Opp complexes together play a significant role in nutrient acquisition and virulence gene activation *in vivo*. Notably, the Δ*oppDF* mutant only had a significant defect in livers of infected mice, though it appears to have a ∼2-fold defect in spleens as well that is not rescued in the PrfA* background, further emphasizing the importance of OppDF in intracellular nutrient acquisition (Figs. 6).

**Figure 6.**
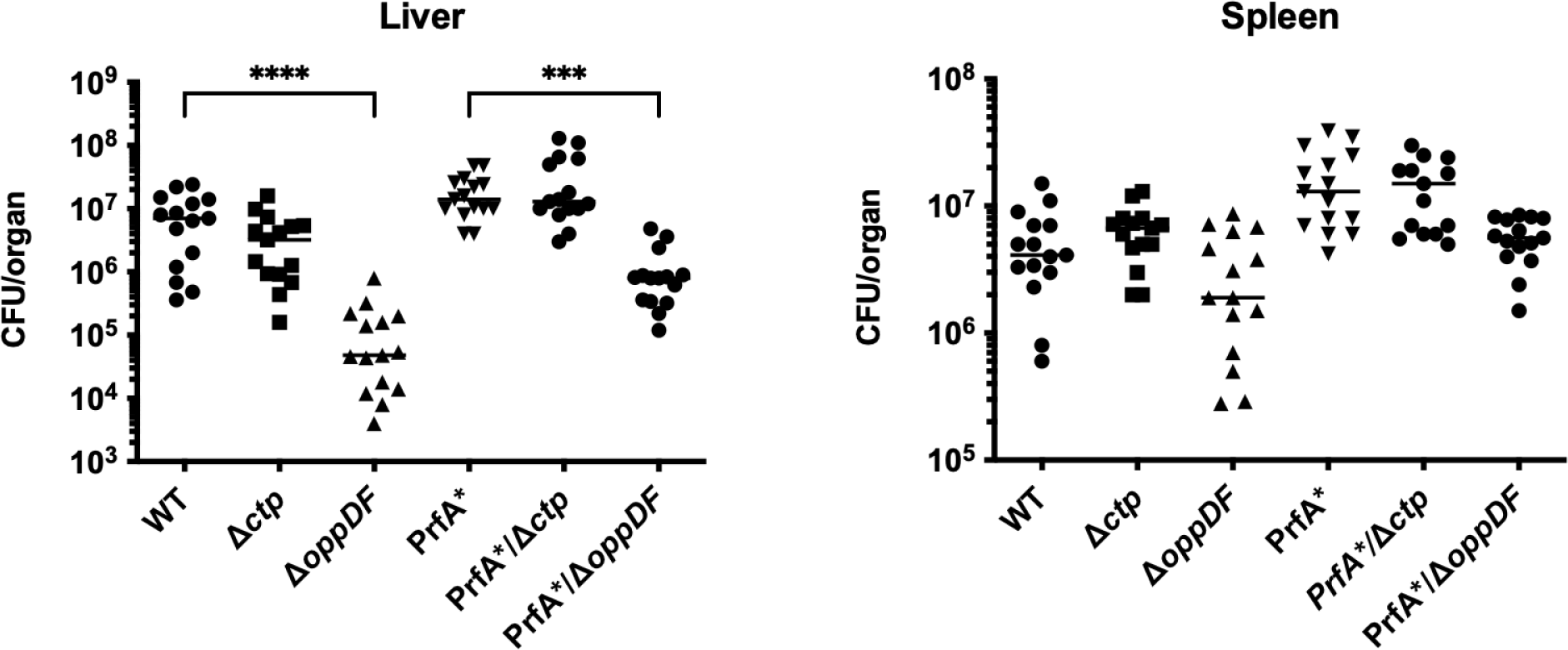
The Ctp/OppDF complex is required for full virulence in a murine IV infection model. 8-week old female CD-1 mice were intravenously infected with 10^5^ CFU of indicated *Lm* strains. Livers and spleens were harvested at 48 hours post infection and bacterial burden was measured by CFU. Data representative of three independent experiments. Statistical significance determined by Kruskal-Wallis test; ***P < 0.0002, ****P < 0.0001.

Overall, our data suggest that the Ctp complex is dispensable for virulence and nutrient acquisition *in vivo* but do not rule out the existence of an alternative redundant GSH importer. The data also suggest that the Ctp and Opp complexes are together required for growth and virulence *in vivo*.

## Discussion

*Listeria monocytogenes* is a pathogen of humans and other mammals that alternates between a saprophytic and intracellular lifecycle, requiring the production and import of GSH to drive virulence gene expression and pathogenesis. Even though L-cysteine is required for GSH synthesis, *Lm* is thought to be auxotrophic for L-cysteine, requiring import of exogenous L-cysteine sources to support growth and virulence (22, 23). Locally high GSH concentrations in the host cytosol have been posited to act as a signal for bacterial entry into the intracellular niche, as the binding affinity of PrfA for GSH has been reported to be ∼4mM, which is well within the biologically relevant concentrations of the host cytoplasm (6, 11). Despite these observations, no GSH importers had been identified in *Lm*. The results of this study demonstrate that the Ctp complex is a high-affinity GSH and GSSG importer in *Lm*, and that *Lm* is not auxotrophic for L-cysteine and can utilize limited inorganic sulfur sources to synthesize L-cysteine and support growth.

The Ctp ABC transporter operon consists of three subunits, a substrate-binding protein and two permeases, but lacks dedicated ATPase domains (Fig. 3A). Our results indicate that the two ATPases from the Opp oligopeptide importer complex, OppDF, are also required for growth on GSH and GSSG independently of their role in the Opp complex (Fig. 7A). Preliminary observations also suggest that the Ctp complex may import short cysteine-containing peptides at lower affinities, which is consistent with observations made of the GshT GSH importer in *Streptococcus mutans* that also facilitates growth on GSH derivatives (27). Despite the observation that the Ctp complex is a high-affinity GSH/GSSG importer, we observed growth of the *ctp* mutant on higher concentrations of exogenous GSH (Fig. 5B), suggesting that an alternative, lower-affinity GSH acquisition mechanism also exists. The most likely candidates for this mechanism are either an additional yet-unidentified GSH importer or a secreted γ-glutamyl transpeptidase (GGT, Fig. 7A), such as that utilized by *Francisella tularensis* (26). The importance of this alternative mechanism(s) to pathogenesis is accentuated by the observation that the *ctp* mutant does not display significant virulence defects in a murine IV infection model, and that the defects in the *oppDF* mutant are restricted to the liver where intracellular GSH concentrations are greatest (Fig. 6, 10). Further work will be required to elucidate any additional GSH acquisition mechanisms.

**Figure 7.**
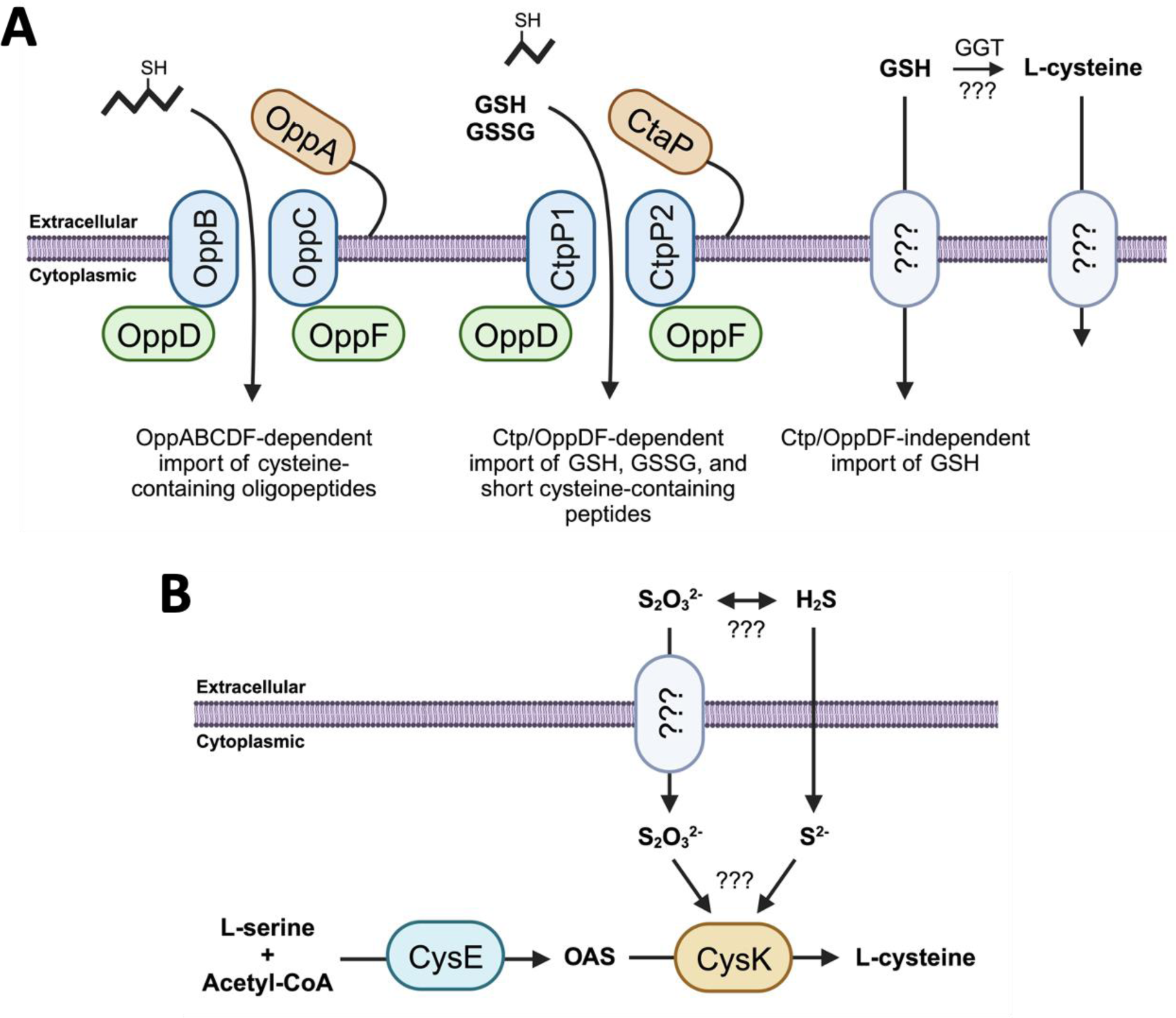
Model of mechanisms for *Lm* growth on GSH, GSSG, thiosulfate, and H_2_S. A) Cysteine-containing oligopeptides are imported by OppABCDF complex. GSH and GSSG are imported by the Ctp/OppDF complex with high affinity, and small cysteine-containing oligopeptides are likely imported with low affinity. An unknown GSH importer and/or a secreted GSH-specific ɣ-glutamyl transpeptidase (GGT) and L-cysteine importer likely also exist that support growth on higher concentrations of GSH. H_2_S freely diffuses through the plasma membrane and/or thiosulfate is imported by an unknown mechanism, where they serve as sulfide donors for production of L-cysteine by the CysE/CysK two-step biosynthetic pathway. It is unclear if H_2_S or thiosulfate is the preferred substrate for CysK. OAS: O-acetylserine, S_2_O_3_^2-^: thiosulfate, H_2_S: hydrogen sulfide.

Another novel finding of this work is that *Lm* utilizes the inorganic sulfur sources thiosulfate and H_2_S as an L-cysteine source. For decades, the consensus was that *Lm* is auxotrophic for L-cysteine despite the presence of an intact and conserved two-step L-cysteine biosynthetic pathway (21–23, 32). Here, we show that the CysE/CysK two-step L-cysteine biosynthetic pathway is functional in *Lm* and both exogenous thiosulfate and H_2_S support growth (Fig. 7B). Because growth is only supported on limited inorganic sulfur sources, our results support a model of partial L-cysteine auxotrophy in *Lm*. Despite our observations, it is not yet clear if or how *Lm* imports thiosulfate, which is a charged molecule that requires facilitated import. Thiosulfate is canonically imported by members of the sulfate assimilation pathway, which is missing in *Lm* (21, 41, 42), however import has also been shown to occur through independent importers and raises the possibility of identifying one or more importers in *Lm* (35, 43). Alternatively, *Lm* could potentially reduce thiosulfate to H_2_S for passive import through the plasma membrane, however this is likely an unfavorable reaction in aerobic environments and little evidence exists to suggest this mechanism.

Another outstanding question is if the CysE/CysK pathway contributes to *Lm* growth and pathogenesis, as concentrations of H_2_S and thiosulfate are uniquely high in the intestinal lumen of mammals. Here, sulfur-reducing bacteria (SRBs) produce millimolar quantities of H_2_S in the anoxic core of the intestine, which is oxidized to thiosulfate at the oxic-anoxic interface generated by oxygen species released by colonic epithelial cells (28–30, 36, 37). Bacterial pathogens are known to exploit these high concentrations of thiosulfate in the gut to facilitate colonization (44), suggesting that *Lm* may have adapted to do so as well. Our results do not indicate that the CysE/CysK pathway alone is required for intestinal colonization in mice (Fig. S2), however it is important to note that the lumen of the intestine is a complex nutrient-rich environment that contains many other exogenous L-cysteine sources. It is also important to consider that streptomycin treatment was used to reduce colonization resistance in the gut, but that the resulting dysbiosis could negatively impact populations of SRBs in the gut and alter the H_2_S/thiosulfate landscape in the intestinal lumen in our experimental model, though little is known about the impacts of antibiotics on SRBs (45). Despite these observations, it is possible that the CysE/CysK pathway still contributes overall L-cysteine levels during infection, and high levels of these inorganic sulfur compounds in the intestine may represent an early host signal that primes *Lm* for intracellular PrfA activation. More work is needed to determine the exact role of the CysE/CysK biosynthetic pathway in the dynamics of host colonization and pathogenesis.

Finally, this study calls into question the very nature of partial L-cysteine auxotrophy in *Listeria monocytogenes*. Auxotrophies constrain the interactions of bacteria with their environment, and bacterial pathogens are no different. While auxotrophies can arise through deleterious mutation, they often arise as specific adaptations to unique host or environmental conditions, suggesting a pathoadaptive role for these auxotrophies (46, 47). *Lm* has a handful of well-characterized auxotrophies that are relevant to pathogenesis, namely lipoic acid and riboflavin. With lipoic acid, host scavenging appears to be essential for intracellular growth, suggesting an adaptation to a unique feature of the host environment (48). In the case of riboflavin, previous work has suggested that WT *Lm* has lost the ability to synthesize a key riboflavin intermediate, 5-OP-RU, to enable evasion of bacterial recognition by host MAIT innate immune cells (49, 50). In both cases, the inability to produce these two essential metabolites confers a fitness advantage to *Lm* in the host environment.

Curiously, L-cysteine auxotrophies in other pathogenic bacterial species (*S. enterica* and *E. coli*) confer a fitness advantage to these bacteria in certain *in vivo* niches (47), suggesting that the same may be true in *Lm*. The pathogen *Staphylococcus aureus* is similarly partially auxotrophic for L-cysteine, lacking an intact sulfate assimilation pathway but can utilize limited inorganic sulfur sources like thiosulfate for growth, and recent work has highlighted the importance of L-cysteine acquisition mechanisms during infection (51–53). Considering the essential role of L-cysteine for growth and virulence gene expression in *Lm*, it is compelling to suggest that partial L-cysteine auxotrophy is a pathoadaptive feature of *Lm* metabolism, representing an adaptation to a unique host environment. Virulence gene activation is tightly regulated in *Lm* and inappropriate activation during saprophytic growth results in considerable defects in bacterial fitness, highlighting the importance of proper spatiotemporal control of PrfA activation (54, 55). Together, these observations suggest that locally high L-cysteine concentrations in the host cytoplasm may serve as a vital host cue that signals bacterial arrival to their intracellular niche. Previous work lends support to this hypothesis, as the peptide signature of the habitat and the relatively higher concentrations of cysteine-containing peptides in the host cytoplasm have been postulated to directly impact GSH production and PrfA activation (13). The host cytoplasm contains uniquely high concentrations of GSH, and this activator of virulence and vital L-cysteine source represent an attractive and specific host signal that bacteria have arrived at their intracellular niche. The discovery of the first GSH-specific importer in *Lm* and the ability to utilize limited inorganic sulfur sources for growth are significant steps in understanding the nature of partial L-cysteine auxotrophy and the spatiotemporal dynamics of virulence gene activation in the host-pathogen interactions of *Listeria monocytogenes*.

## Materials and Methods

### Bacterial strains, plasmids, and growth conditions

All strains of *L. monocytogenes* used in this study were derived from the WT 10403S parental strain. Strains were cultured in chemically defined synthetic media (LSM) containing 200ug/mL streptomycin (Sigma-Aldrich) and 0.5mM L-cystine (Sigma-Aldrich) as the sole cysteine source. LSM was prepared as previously described (56). LSM containing the alternative cysteine sources reduced L-cysteine (Sigma-Aldrich), reduced glutathione (Sigma-Aldrich), oxidized glutathione (Sigma-Aldrich), RGDC oligopeptide (GenScript), sodium thiosulfate (Sigma-Aldrich), and sodium hydrosulfide (Sigma-Aldrich) was prepared by substituting L-cystine for the indicated concentration of each cysteine source. Media containing reduced L-cysteine or reduced glutathione were prepared fresh and supplemented with 2mM TCEP (Goldbio) prior to experimentation, except where otherwise stated. Media containing sodium hydrosulfide was prepared fresh immediately prior to experimentation to prevent off-gassing of volatile hydrogen sulfide gas. Additional antibiotics (Sigma-Aldrich) were used at the following concentrations: erythromycin (1ug/mL), carbenicillin (100ug/mL), tetracycline (2ug/mL), and chloramphenicol (7.5ug/mL for *L. monocytogenes* and 10ug/mL for *E. coli*).

Construction of in-frame deletion strains Δ*ctp*, Δ*ctpP1*, Δ*oppB*, Δ*oppDF*, and Δ*c*ysK was performed by allelic exchange using the temperature-sensitive vector pKSV7 as previously described (57). Nutrient-rich Brain Heart Infusion media (BHI, BD Difco) was used for bacterial growth during the allelic exchange protocol. To circumvent growth deficiencies of the Δ*ctp*, Δ*ctpP1*, and Δ*oppDF* mutants in BHI, mutants were grown in BHI supplemented with all 20 amino acids (Sigma-Aldrich) as in LSM media (0.1g/L, except L-glutamine at 0.6g/L).

Complementation of genes was performed by amplifying the native genes from WT 10403S *L. monocytogenes* and cloning into the site-specific pPL2 integrating vector containing the constitutively expressed pHyper promoter. Complementation vectors were conjugated into *L. monocytogenes* as previously described (58).

### Broth growth assays

Broth growth assays were performed with *L. monocytogenes* strains grown overnight in LSM supplemented with 0.5mM L-cystine at 37°C with shaking (220rpm). Bacteria were washed once in synthetic media lacking an L-cysteine source (LSM-cys) and resuspended in an equal volume of LSM-cys. Washed bacteria were diluted to an optical density of 0.05 in LSM-cys containing the indicated cysteine sources and grown at 37°C with shaking. Growth was assessed spectrophotometrically by optical density at a wavelength of 600nm (OD_600_). Broth growth curves were performed in 250mL flasks containing 35mL of media and OD_600_ measurements were taken every two hours for 14 hours and again at 24-26 hours as indicated. Saturation endpoint growth experiments were performed in 14mL round-bottom tubes containing 2mL of media and OD_600_ measurements were taken at 24-26 hours post-inoculation.

### Mouse intravenous infections

Eight-week-old female CD-1 mice (Charles River Laboratories) were infected intravenously via the tail vein with 200uL PBS containing 1×10^5^ CFU of logarithmically growing *L. monocytogenes* as previously described (59). Bacteria used for infection were grown overnight in LSM supplemented with 0.5mM L-cystine and diluted 1:20 in fresh media to obtain logarithmically growing bacteria. Mice were euthanized 48 hours post-infection and spleens, livers, and gallbladders were collected, homogenized, and plated to enumerate CFUs per organ. Statistical significance was determined by one-way ANOVA and Kruskal-Wallis test.

### Mouse oral infection competition assay

Eight-week-old female C57BL/6J mice (Jackson Laboratories) were provided 5ug/mL streptomycin (Sigma-Aldrich) in drinking water 48 hours prior to infection as previously described (60). Mice were transferred to clean cages 18-24 hours prior to infection and the food source was removed to fast mice overnight. Bacteria used for infection (Δ*hly* and Δ*hly*/ErmR) were grown overnight in BHI and diluted 1:10 in fresh media to obtain logarithmically growing bacteria. At time of infection, bacteria were washed twice in PBS and inoculums containing 1×10^8^ each of Δ*hly* and Δ*hly*/ErmR (WT control or mutant) *L. monocytogenes* were prepared in 5uL PBS. A 3-mm piece of bread was inoculated with a total of 2×10^8^ CFU bacteria in PBS and covered with 3uL melted butter. Mice were fed a single piece of infected bread. Streptomycin-treated water was replaced with standard drinking water and food was restored at the time of inoculation. Fecal samples were collected every day post-infection for five days, weighed, and resuspended in 1mL PBS by vortexing at 4°C for 10 minutes. Fecal samples were plated on BHI plates containing only streptomycin or streptomycin and erythromycin to enumerate CFUs of WT and erythromycin-resistant strains and the ratio of the two populations was calculated to determine the competitive fitness of mutant strains relative to WT 10403S.

### Animal Use Ethics Statement

Mice were maintained by University of California, Berkeley Office of Laboratory Animal Care personnel according to institutional guidelines. All animal studies were performed in accordance with the recommendation in the Guide for the Care and Use of Laboratory Animals of the National Institutes of Health and university regulations. The protocols used in this study were reviewed and approved by the Animal Care and Use Committee at the University of California, Berkeley (AUP 2016-05-8811).

### Statistical analysis

All statistical analysis was performed using GraphPad Prism version 10.

## Acknowledgements

The research reported in this publication was supported by National Institutes of Health grants 1P01AI063302 and 1R01AI27655 to D.A.P, and by the National Science Foundation Graduate Research Fellowship grant DGE 1752814 to J.C.B. P.K. was supported by the Département de Biologie, École Normale Supérieure de Lyon, Lyon, France. The funders had no role in study design, data collection and analysis, decision to publish, or preparation of the manuscript. D.A.P. has a financial interest in Laguna Biotherapeutics, and both he and the company could benefit from the commercialization of the results of this research. The models presented in Figures 2A, 2B, 3A, and 7 were created with BioRender.com.

---

**Figure S1.**
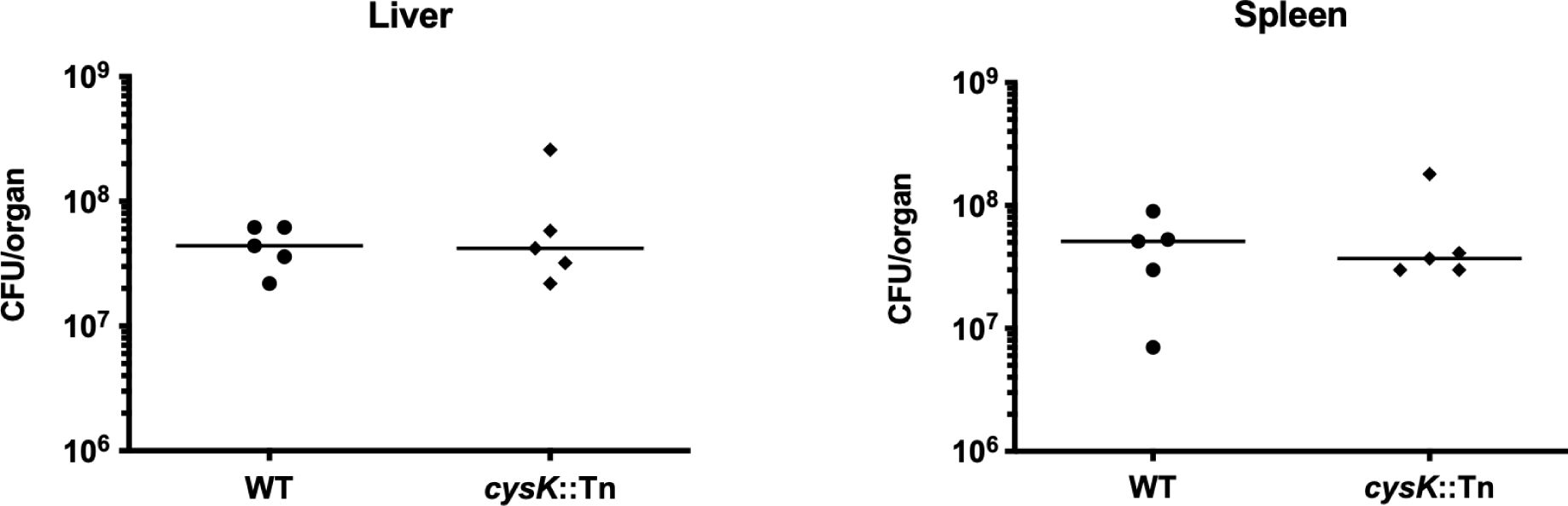
CysK is not required for virulence in a murine IV infection model. 8-week old female CD-1 mice were intravenously infected with 10^5^ CFU of WT or *cysK*::Tn mutant *Lm*. Livers and spleens were harvested at 48 hours post infection and bacterial burden was measured by CFU. Data representative of one replicate.

**Figure S2.**
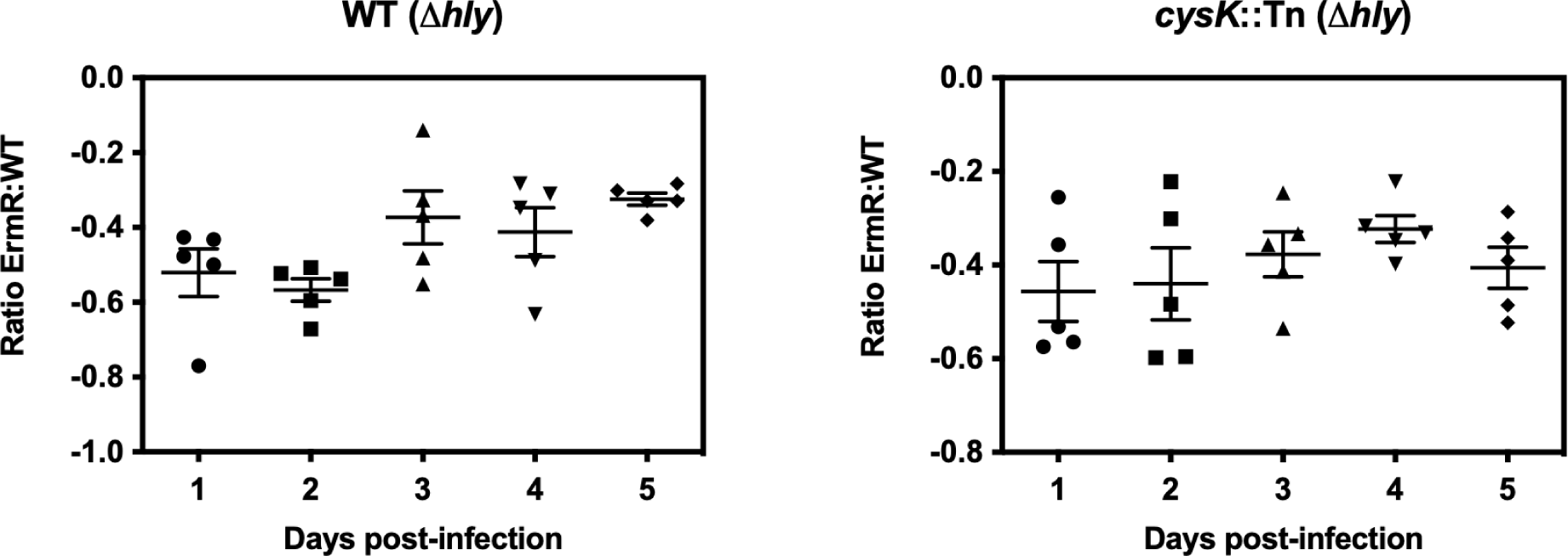
CysK is not required for bacterial fitness in a murine oral infection model. Streptomycin-treated 8-week old female C57BL/6J mice were orally infected with 10^8^ CFU each of *Δhly* and *Δhly* containing the Erm resistance gene. Fecal pellets were collected every day for five days post-infection, CFUs were enumerated for total and Erm resistant bacterial populations, and the ratio of ErmR:WT bacteria was calculated. Data representative of one replicate.

**Figure S3.**
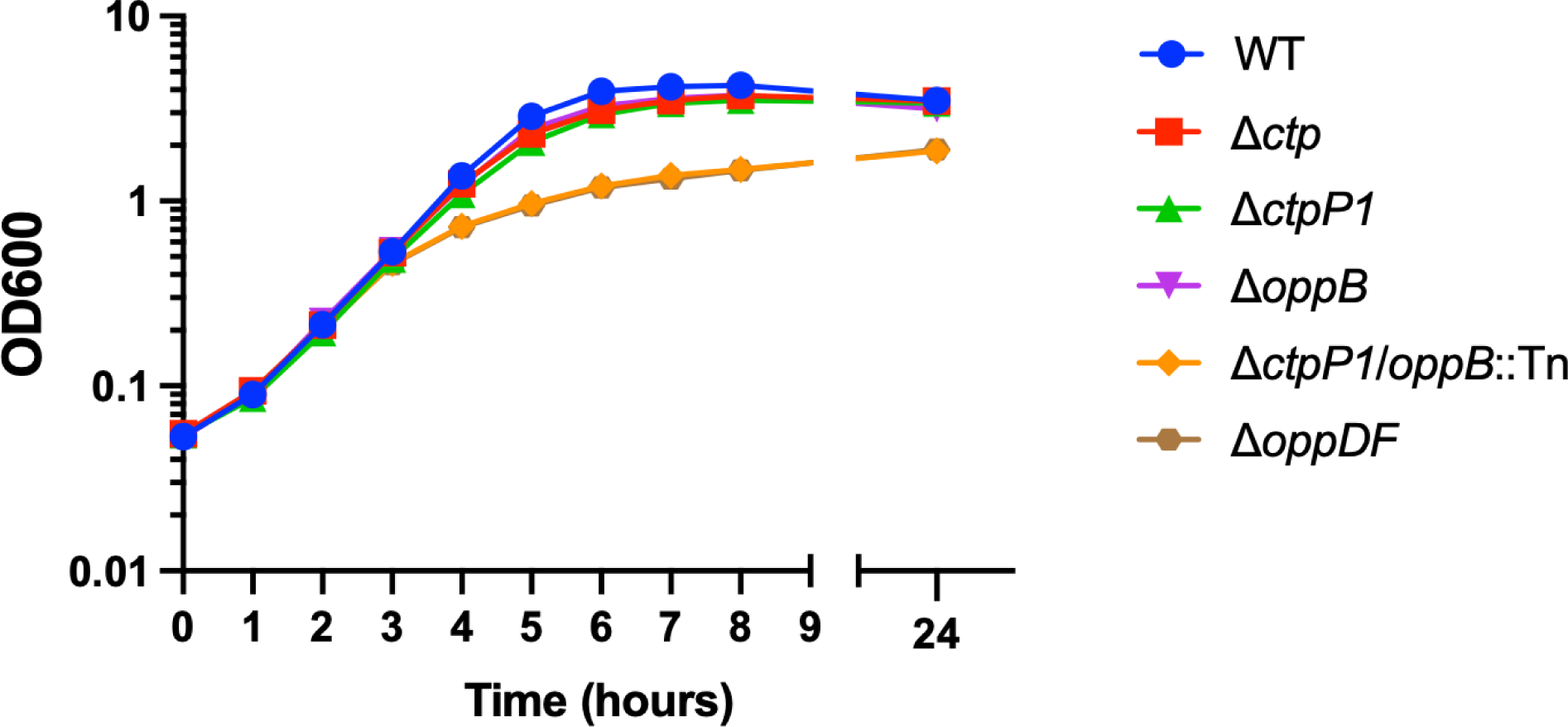
OppDF and the Ctp/Opp double mutant demonstrate growth defects in complex rich media. Broth growth of the *ctp* and *opp* mutants in Brain-Heart Infusion (BHI) medium. Strains were grown overnight in LSM media containing 0.5mM L-cystine, washed once in BHI, and inoculated into BHI. Growth was determined by OD_600_ at the indicated timepoints. Data representative of three independent replicates, error bars represent SD.

**Table S1.**
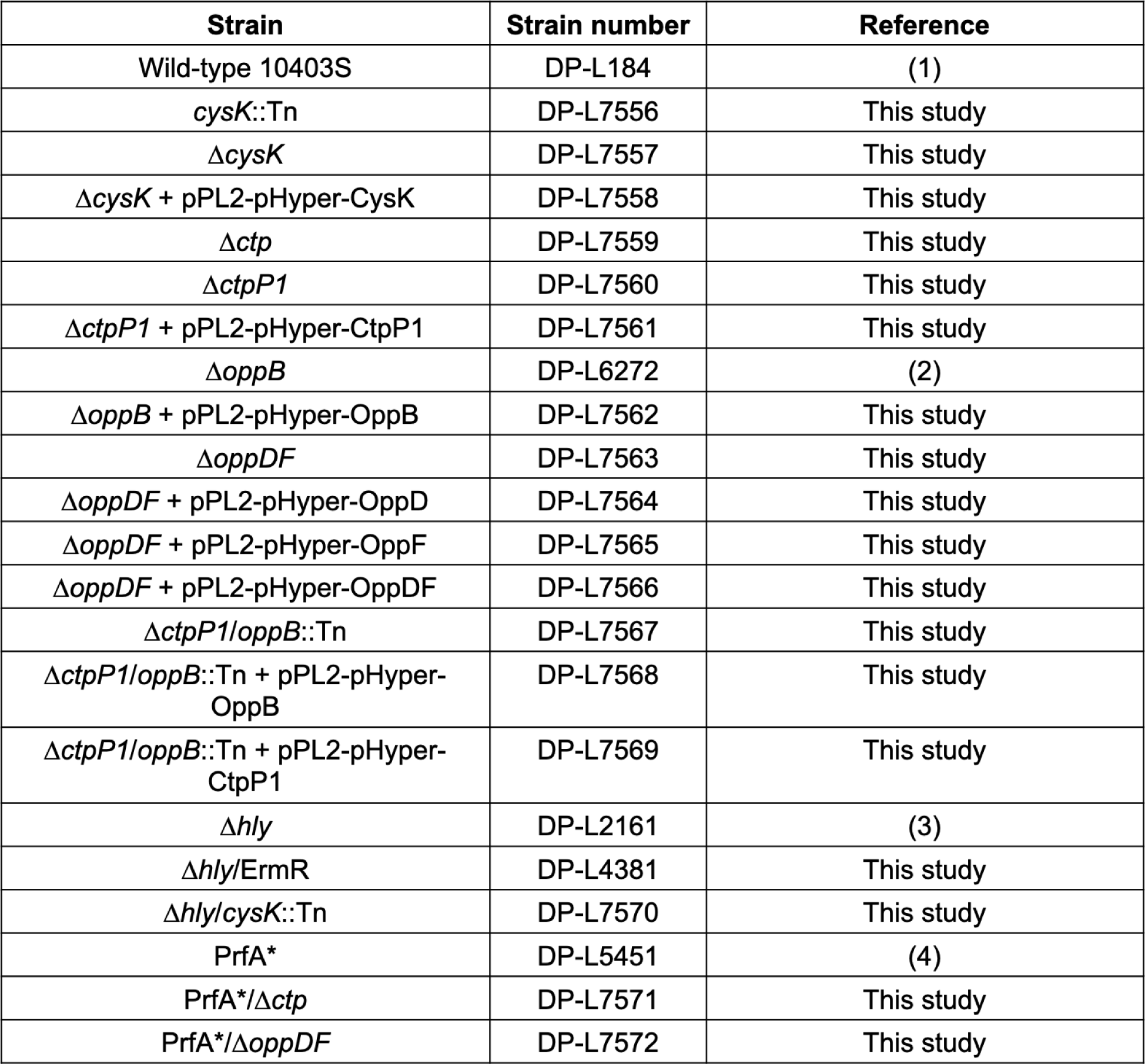
*Listeria monocytogenes* strains used in this study.

**Table S2.** *Escherichia coli* strains used in this study.

| Strain | Strain number | Reference |
| --- | --- | --- |
| SM10 pKSV7x-*cysK* | DP-L7573 | This study |
| SM10 pKSV7x-*ctp* | DP-L7574 | This study |
| SM10 pKSV7x-*ctpP1* | DP-L7575 | This study |
| SM10 pKSV7x-*oppDF* | DP-L7576 | This study |
| SM10 pPL2x-pHyper-CysK | DP-L7577 | This study |
| SM10 pPL2x-pHyper-CtpP1 | DP-L7578 | This study |
| SM10 pPL2x-pHyper-OppB | DP-L7579 | This study |
| SM10 pPL2x-pHyper-OppD | DP-L7580 | This study |
| SM10 pPL2x-pHyper-OppF | DP-L7581 | This study |
| SM10 pPL2x-pHyper-OppDF | DP-L7582 | This study |

**Table S3.**
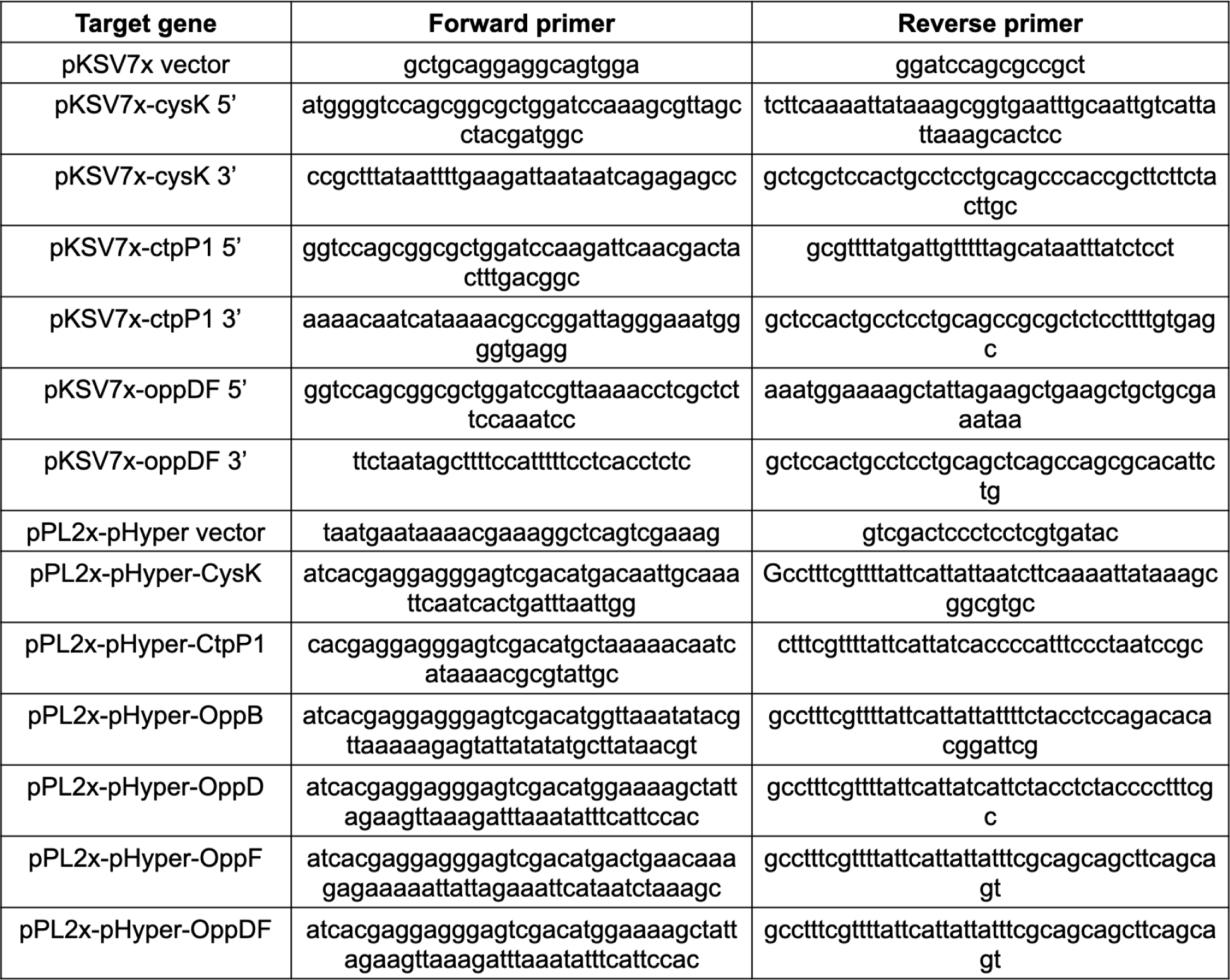
Oligonucleotides strains used in this study.

## References

1. Freitag NE, Port GC, Miner MD. 2009. Listeria monocytogenes - from saprophyte to intracellular pathogen. Nat Rev Microbiol 7:623–8.

2. Koopmans MM, Brouwer MC, Vázquez-Boland JA, van de Beek D. 2023. Human Listeriosis. Clin Microbiol Rev 36:e0006019.

3. Freitag NE. 2006. From hot dogs to host cells: how the bacterial pathogen Listeria monocytogenes regulates virulence gene expression. Future Microbiol 1:89–101.

4. Swaminathan B, Gerner-Smidt P. 2007. The epidemiology of human listeriosis. Microbes Infect 9:1236–43.

5. de las Heras A, Cain RJ, Bielecka MK, Vázquez-Boland JA. 2011. Regulation of Listeria virulence: PrfA master and commander. Curr Opin Microbiol 14:118–27.

6. Reniere ML, Whiteley AT, Hamilton KL, John SM, Lauer P, Brennan RG, Portnoy DA. 2015. Glutathione activates virulence gene expression of an intracellular pathogen. Nature 517:170–3.

7. Hall M, Grundström C, Begum A, Lindberg MJ, Sauer UH, Almqvist F, Johansson J, Sauer-Eriksson AE. 2016. Structural basis for glutathione-mediated activation of the virulence regulatory protein PrfA in Listeria. Proc Natl Acad Sci U S A 113:14733–14738.

8. Anaya-Sanchez A, Feng Y, Berude JC, Portnoy DA. 2021. Detoxification of methylglyoxal by the glyoxalase system is required for glutathione availability and virulence activation in Listeria monocytogenes. PLoS Pathog 17:e1009819.

9. Masip L, Veeravalli K, Georgiou G. 2006. The many faces of glutathione in bacteria. Antioxid Redox Signal 8:753–62.

10. Meister A, Anderson ME. 1983. Glutathione. Annu Rev Biochem 52:711–60.

11. Portman JL, Dubensky SB, Peterson BN, Whiteley AT, Portnoy DA. 2017. Activation of the Listeria monocytogenes virulence program by a reducing environment. mBio 8.

12. Brenner M, Friedman S, Haber A, Livnat-Levanon N, Borovok I, Sigal N, Lewinson O, Herskovits AA. 2022. Listeria monocytogenes TcyKLMN Cystine/Cysteine Transporter Facilitates Glutathione Synthesis and Virulence Gene Expression. mBio 13:e0044822.

13. Krypotou E, Scortti M, Grundström C, Oelker M, Luisi BF, Sauer-Eriksson AE, Vázquez-Boland J. 2019. Control of Bacterial Virulence through the Peptide Signature of the Habitat. Cell Rep 26:1815–1827.e5.

14. Park S, Imlay JA. 2003. High levels of intracellular cysteine promote oxidative DNA damage by driving the fenton reaction. J Bacteriol 185:1942–50.

15. Korshunov S, Imlay KRC, Imlay JA. 2020. Cystine import is a valuable but risky process whose hazards Escherichia coli minimizes by inducing a cysteine exporter. Mol Microbiol 113:22–39.

16. Zhou Y, Imlay JA. 2020. Escherichia coli K-12 Lacks a High-Affinity Assimilatory Cysteine Importer. mBio 11.

17. Zhou Y, Imlay JA. 2022. Escherichia coli Uses a Dedicated Importer and Desulfidase To Ferment Cysteine. mBio 13:e0296521.

18. Xayarath B, Marquis H, Port GC, Freitag NE. 2009. Listeria monocytogenes CtaP is a multifunctional cysteine transport-associated protein required for bacterial pathogenesis. Mol Microbiol 74:956–73.

19. Vaval Taylor DM, Xayarath B, Freitag NE. 2023. Two Permeases Associated with the Multifunctional CtaP Cysteine Transport System in Listeria monocytogenes Play Distinct Roles in Pathogenesis. Microbiol Spectr 11:e0331722.

20. Fujishima K, Wang KM, Palmer JA, Abe N, Nakahigashi K, Endy D, Rothschild LJ. 2018. Reconstruction of cysteine biosynthesis using engineered cysteine-free enzymes. Sci Rep 8:1776.

21. Glaser P, Frangeul L, Buchrieser C, Rusniok C, Amend A, Baquero F, Berche P, Bloecker H, Brandt P, Chakraborty T, Charbit A, Chetouani F, Couvé E, de Daruvar A, Dehoux P, Domann E, Domínguez-Bernal G, Duchaud E, Durant L, Dussurget O, Entian KD, Fsihi H, García-del Portillo F, Garrido P, Gautier L, Goebel W, Gómez-López N, Hain T, Hauf J, Jackson D, Jones LM, Kaerst U, Kreft J, Kuhn M, Kunst F, Kurapkat G, Madueno E, Maitournam A, Vicente JM, Ng E, Nedjari H, Nordsiek G, Novella S, de Pablos B, Pérez-Diaz JC, Purcell R, Remmel B, Rose M, Schlueter T, Simoes N, et al. 2001. Comparative genomics of Listeria species. Science 294:849–52.

22. Tsai HN, Hodgson DA. 2003. Development of a synthetic minimal medium for Listeria monocytogenes. Appl Environ Microbiol 69:6943–5.

23. Sauer JD, Herskovits AA, O’Riordan MXD. 2019. Metabolism of the Gram-Positive Bacterial Pathogen Listeria monocytogenes. Microbiol Spectr 7.

24. Parry J, Clark DP. 2002. Identification of a CysB-regulated gene involved in glutathione transport in Escherichia coli. FEMS Microbiol Lett 209:81–5.

25. Suzuki H, Koyanagi T, Izuka S, Onishi A, Kumagai H. 2005. The yliA, -B, -C, and -D genes of Escherichia coli K-12 encode a novel glutathione importer with an ATP-binding cassette. J Bacteriol 187:5861–7.

26. Alkhuder K, Meibom KL, Dubail I, Dupuis M, Charbit A. 2009. Glutathione provides a source of cysteine essential for intracellular multiplication of Francisella tularensis. PLoS Pathog 5:e1000284.

27. Vergauwen B, Verstraete K, Senadheera DB, Dansercoer A, Cvitkovitch DG, Guédon E, Savvides SN. 2013. Molecular and structural basis of glutathione import in Gram-positive bacteria via GshT and the cystine ABC importer TcyBC of Streptococcus mutans. Mol Microbiol 89:288–303.

28. Singh P, Teal TK, Marsh TL, Tiedje JM, Mosci R, Jernigan K, Zell A, Newton DW, Salimnia H, Lephart P, Sundin D, Khalife W, Britton RA, Rudrik JT, Manning SD. 2015. Intestinal microbial communities associated with acute enteric infections and disease recovery. Microbiome 3:45.

29. Imlay JA. 2019. Where in the world do bacteria experience oxidative stress? Environ Microbiol 21:521–530.

30. Dordević D, Jančíková S, Vítězová M, Kushkevych I. 2021. Hydrogen sulfide toxicity in the gut environment: Meta-analysis of sulfate-reducing and lactic acid bacteria in inflammatory processes. Journal of Advanced Research 27:55–69.

31. Nepal S, Kumar P. 2020. Growth, Cell Division, and Gene Expression of Escherichia coli at Elevated Concentrations of Magnesium Sulfate: Implications for Habitability of Europa and Mars. Microorganisms 8.

32. Premaratne RJ, Lin WJ, Johnson EA. 1991. Development of an improved chemically defined minimal medium for Listeria monocytogenes. Appl Environ Microbiol 57:3046–8.

33. Kawano Y, Suzuki K, Ohtsu I. 2018. Current understanding of sulfur assimilation metabolism to biosynthesize L-cysteine and recent progress of its fermentative overproduction in microorganisms. Appl Microbiol Biotechnol 102:8203–8211.

34. Mathai JC, Missner A, Kügler P, Saparov SM, Zeidel ML, Lee JK, Pohl P. 2009. No facilitator required for membrane transport of hydrogen sulfide. Proc Natl Acad Sci U S A 106:16633–8.

35. Tanaka Y, Yoshikaie K, Takeuchi A, Ichikawa M, Mori T, Uchino S, Sugano Y, Hakoshima T, Takagi H, Nonaka G, Tsukazaki T. 2020. Crystal structure of a YeeE/YedE family protein engaged in thiosulfate uptake. Sci Adv 6:eaba7637.

36. Furne J, Saeed A, Levitt MD. 2008. Whole tissue hydrogen sulfide concentrations are orders of magnitude lower than presently accepted values. Am J Physiol Regul Integr Comp Physiol 295:R1479–85.

37. Karunya R, Jayaprakash KS, Gaikwad R, Sajeesh P, Ramshad K, Muraleedharan KM, Dixit M, Thangaraj PR, Sen AK. 2019. Rapid measurement of hydrogen sulphide in human blood plasma using a microfluidic method. Sci Rep 9:3258.

38. Tomich PK, An FY, Clewell DB. 1980. Properties of erythromycin-inducible transposon Tn917 in Streptococcus faecalis. J Bacteriol 141:1366–74.

39. Borezee E, Pellegrini E, Berche P. 2000. OppA of Listeria monocytogenes, an oligopeptide-binding protein required for bacterial growth at low temperature and involved in intracellular survival. Infect Immun 68:7069–77.

40. Ripio MT, Domínguez-Bernal G, Lara M, Suárez M, Vazquez-Boland JA. 1997. A Gly145Ser substitution in the transcriptional activator PrfA causes constitutive overexpression of virulence factors in Listeria monocytogenes. J Bacteriol 179:1533–40.

41. Sirko A, Zatyka M, Sadowy E, Hulanicka D. 1995. Sulfate and thiosulfate transport in Escherichia coli K-12: evidence for a functional overlapping of sulfate- and thiosulfate-binding proteins. J Bacteriol 177:4134–6.

42. Hicks JL, Mullholland CV. 2018. Cysteine biosynthesis in Neisseria species. Microbiology (Reading) 164:1471–1480.

43. Mansilla MC, de Mendoza D. 2000. The Bacillus subtilis cysP gene encodes a novel sulphate permease related to the inorganic phosphate transporter (Pit) family. Microbiology (Reading) 146 (Pt 4):815–821.

44. Winter SE, Thiennimitr P, Winter MG, Butler BP, Huseby DL, Crawford RW, Russell JM, Bevins CL, Adams LG, Tsolis RM, Roth JR, Bäumler AJ. 2010. Gut inflammation provides a respiratory electron acceptor for Salmonella. Nature 467:426–9.

45. Karnachuk OV, Beletsky AV, Rakitin AL, Ikkert OP, Avakyan MR, Zyusman VS, Napilov A, Mardanov AV, Ravin NV. 2023. Antibiotic-resistant Desulfovibrio produces H_2_S from supplements for animal farming. Microorganisms 11.

46. Sokurenko EV, Hasty DL, Dykhuizen DE. 1999. Pathoadaptive mutations: gene loss and variation in bacterial pathogens. Trends Microbiol 7:191–5.

47. Seif Y, Choudhary KS, Hefner Y, Anand A, Yang L, Palsson BO. 2020. Metabolic and genetic basis for auxotrophies in Gram-negative species. Proc Natl Acad Sci U S A 117:6264–6273.

48. O’Riordan M, Moors MA, Portnoy DA. 2003. Listeria intracellular growth and virulence require host-derived lipoic acid. Science 302:462–4.

49. Legoux F, Salou M, Lantz O. 2020. MAIT Cell Development and Functions: the Microbial Connection. Immunity 53:710–723.

50. Rivera-Lugo R, Light SH, Garelis NE, Portnoy DA. 2022. RibU is an essential determinant of Listeria pathogenesis that mediates acquisition of FMN and FAD during intracellular growth. Proc Natl Acad Sci U S A 119:e2122173119.

51. Lithgow JK, Hayhurst EJ, Cohen G, Aharonowitz Y, Foster SJ. 2004. Role of a cysteine synthase in Staphylococcus aureus. J Bacteriol 186:1579–90.

52. Lensmire JM, Dodson JP, Hsueh BY, Wischer MR, Delekta PC, Shook JC, Ottosen EN, Kies PJ, Ravi J, Hammer ND. 2020. The Staphylococcus aureus Cystine Transporters TcyABC and TcyP Facilitate Nutrient Sulfur Acquisition during Infection. Infect Immun 88.

53. Lensmire JM, Wischer MR, Kraemer-Zimpel C, Kies PJ, Sosinski L, Ensink E, Dodson JP, Shook JC, Delekta PC, Cooper CC, Havlichek DH, Mulks MH, Lunt SY, Ravi J, Hammer ND. 2023. The glutathione import system satisfies the Staphylococcus aureus nutrient sulfur requirement and promotes interspecies competition. PLoS Genet 19:e1010834.

54. Bruno JC, Freitag NE. 2010. Constitutive activation of PrfA tilts the balance of Listeria monocytogenes fitness towards life within the host versus environmental survival. PLoS One 5:e15138.

55. Vasanthakrishnan RB, de Las Heras A, Scortti M, Deshayes C, Colegrave N, Vázquez-Boland JA. 2015. PrfA regulation offsets the cost of Listeria virulence outside the host. Environ Microbiol 17:4566–79.

56. Whiteley AT, Garelis NE, Peterson BN, Choi PH, Tong L, Woodward JJ, Portnoy DA. 2017. c-di-AMP modulates Listeria monocytogenes central metabolism to regulate growth, antibiotic resistance and osmoregulation. Mol Microbiol 104:212–233.

57. Camilli A, Tilney LG, Portnoy DA. 1993. Dual roles of plcA in Listeria monocytogenes pathogenesis. Mol Microbiol 8:143–57.

58. Lauer P, Chow MY, Loessner MJ, Portnoy DA, Calendar R. 2002. Construction, characterization, and use of two Listeria monocytogenes site-specific phage integration vectors. J Bacteriol 184:4177–86.

59. Rae CS, Geissler A, Adamson PC, Portnoy DA. 2011. Mutations of the Listeria monocytogenes peptidoglycan N-deacetylase and O-acetylase result in enhanced lysozyme sensitivity, bacteriolysis, and hyperinduction of innate immune pathways. Infect Immun 79:3596–606.

60. Louie A, Zhang T, Becattini S, Waldor MK, Portnoy DA. 2019. A Multiorgan Trafficking Circuit Provides Purifying Selection of Listeria monocytogenes Virulence Genes. mBio 10.

## References

1. Bécavin C, Bouchier C, Lechat P, Archambaud C, Creno S, Gouin E, Wu Z, Kühbacher A, Brisse S, Pucciarelli MG, García-del Portillo F, Hain T, Portnoy DA, Chakraborty T, Lecuit M, Pizarro-Cerdá J, Moszer I, Bierne H, Cossart P. 2014. Comparison of widely used Listeria monocytogenes strains EGD, 10403S, and EGD-e highlights genomic variations underlying differences in pathogenicity. mBio 5:e00969–14.

2. Whiteley AT, Pollock AJ, Portnoy DA. 2015. The PAMP c-di-AMP Is Essential for Listeria monocytogenes Growth in Rich but Not Minimal Media due to a Toxic Increase in (p)ppGpp. [corrected]. Cell Host Microbe 17:788–98.

3. Jones S, Portnoy DA. 1994. Characterization of Listeria monocytogenes pathogenesis in a strain expressing perfringolysin O in place of listeriolysin O. Infect Immun 62:5608–13.

4. Miner MD, Port GC, Freitag NE. 2008. Functional impact of mutational activation on the Listeria monocytogenes central virulence regulator PrfA. Microbiology (Reading) 154:3579–3589.

